# Pulmonary Ventilation Analysis Using ^1^H Ultra-Short Echo Time (UTE) Lung MRI: A Reproducibility Study

**DOI:** 10.1101/2023.10.22.563196

**Authors:** Fei Tan, Xucheng Zhu, Marilynn Chan, Nikhil Deveshwar, Matthew M. Willmering, Michael Lustig, Peder E. Z. Larson

## Abstract

**Purpose:** To evaluate methods for quantification of pulmonary ventilation with ultrashort echo time (UTE) MRI.

**Methods:** We performed a reproducibility study, acquiring two free-breathing ^1^H UTE lung MRIs on the same day for six healthy volunteers. The 1) 3D + t cyclic b-spline and 2) symmetric image normalization (SyN) methods for image registration were applied after respiratory phase-resolved image reconstruction. Ventilation maps were calculated using 1) Jacobian determinant of the deformation fields minus one, termed regional ventilation, and 2) intensity percentage difference between the registered and fixed image, termed specific ventilation. We compared the reproducibility of all four method combinations via statistical analysis.

**Results:** Split violin plots and Bland-Altman plots are shown for whole lungs and lung sections. The cyclic b-spline registration and Jacobian determinant regional ventilation quantification provide total ventilation volumes that match the segmentation tidal volume, smooth and uniform ventilation maps. The cyclic b-spline registration and specific ventilation combination yields the smallest standard deviation in the Bland-Altman plot.

**Conclusion:** Cyclic registration performs better than SyN for respiratory phase-resolved ^1^H UTE MRI ventilation quantification. Regional ventilation correlates better with segmentation lung volume, while specific ventilation is more reproducible.

## Introduction

^1^H MRI has recently been explored as a radiation-free alternative for lung ventilation evaluation (1,2). The Fourier decomposition method (3) acquires consecutive 2D MRIs, applies Fourier decomposition along the time dimension at each pixel, and then separates the cardiac motion and pulmonary motion by the frequency spectrum. Phase-resolved functional lung imaging (PREFUL) (4) reorders the images according to the phase of the cardiac or respiratory motion and reconstructs one full cardiac/respiratory cycle. The flow-volume loop (5) regional.

UTE MRI is favorable for free-breathing lung imaging (6,7) because it is robust to motion artifacts and improves signal-to-noise ratio (SNR) from low T2* tissues such as the lung parenchyma (8). The center of k-space can serve as a self-navigator that tracks respiratory motion during data acquisition (9,10). By grouping the k-space data acquired at different respiratory states, we can reconstruct lung images at these states with minimal motion artifacts, which are the basis of ventilation analysis. Examples include XD-GRASP (10) and self-navigated motion-resolved reconstruction (9).

In this work, we evaluated methods for lung ventilation quantification with a free-breathing respiratory phase-resolved 3D radial UTE lung MRI. We evaluated the effects of two registration methods and two ventilation quantification methods by a reproducibility study in healthy volunteers.

## Methods

### Study Design

All procedures were approved by the University of California, San Francisco’s Institutional Review Board (IRB). 6 healthy volunteers (age 23 - 30, 3 Female, 3 Male) were recruited to the study and written consent was acquired before their scan. Eligibility criteria were no history of asthma or smoking.

An efficiency and SNR optimized variable-density 3D radial UTE sequence (11) with golden angle ordering was used on a 3T clinical scanner (Discovery MR750, GE Healthcare, Waukesha, WI) with an 8-channel cardiac phased-array coil (GE Healthcare, Waukesha, WI). Volunteers performed tidal breathing in a supine position. The key scan parameters were: flip angle=4°, FOV=40cm isotropic, TE/TR=0.1/2.4ms, BW=±125kHz, readout points=512, spokes=200,000, resolution=2.5mm isotropic (Figure 1A). For studying reproducibility, we scanned each subject with the same sequences twice in the same day, 30 minutes apart, re-setting up the coils between the scans. The pipeline for the study is shown in Figure 1.

**Figure 1.**
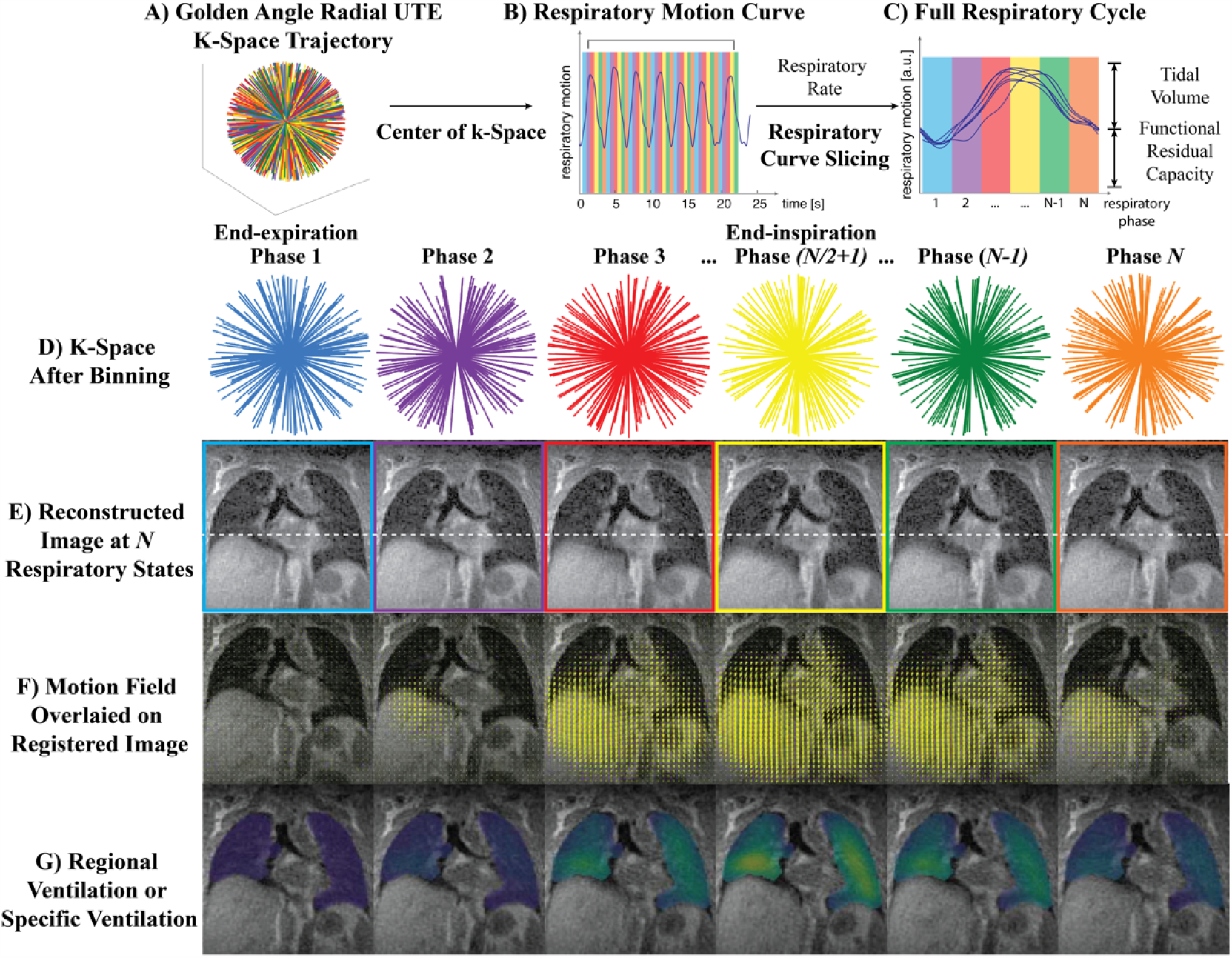
Pipeline for Ventilation Analysis of UTE Lung MRI with N Respiratory Phases. Note that the same procedure is performed twice on the same day for reproducibility studies. A) Data acquisition. B-E) Image reconstruction, the color coding represents respiratory phases. F) Image registration, three types of image registration methods were experimented. G) Ventilation analysis, the Jacobian determinant of the motion field vector at each voxel minus 1, represents the regional ventilation percentage.

### Image Processing

#### Reconstruction

The reconstruction was completed by MATLAB (Mathworks 2019a) and the BART (Berkeley Advanced Reconstruction Toolbox) (12). The data were first binned into 12 phases according to their respiratory cycle timing (Figure 1B, C, D) derived from the center of k-space (9). The number of phases was chosen to balance between less streaking artifacts and more respiratory phases. The respiratory motion was extracted from the coil with the strongest signal fluctuation around the breathing frequency. A 4D phase-resolved image series with three spatial dimensions plus a respiratory phase dimension were reconstructed by parallel imaging and compressed sensing (PICS) reconstruction (Figure 1E).

#### Registration

Then, we registered the image of each respiratory state to the end-expiration state (Figure 1F) with two different methods: 1) B-spline 3D+t cyclic registration (13) to utilize the cyclic characteristic of breathing pattern, and 2) the 3D symmetric image normalization (SyN) method (14) with mutual information metric to minimize the effect of intensity change during breathing. The registrations were implemented in Elastix (15) and ANTs (16) toolboxes, respectively. Both methods utilized 4-level multiscale registration.

#### Segmentation

Segmentation of the left and right lung was achieved by a pre-trained U-Net deep learning algorithm from (17).

### Ventilation Analysis

#### Regional Ventilation

Regional ventilation was calculated based on the local lung tissue motion and deformation information. The Jacobian determinant (JD) of the registration motion field (D) represents the ratio of volume at each respiratory state, *v*_*resp*_, to the end-expiration state, *v*_*end-expi*_. JD >1 means expansion and JD<1 means contraction. The definition was as follows (18,19),

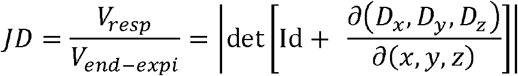

where x,y,z were the spatial dimensions and Id was a 3-by-3 identity matrix.

The regional ventilation definition from (20) was adopted to show the local percentage lung volume change by subtracting 1:

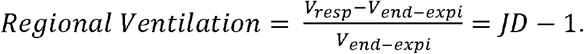

#### Specific Ventilation

Specific ventilation (21,22) was calculated by the percentage intensity difference at each respiratory state to the end-expiration state, measured using the registered images. The definition was as follows,

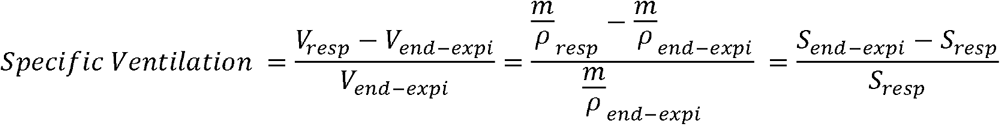

where *S*_*end-expi*_, *S*_*resp*_ were the signal intensities of each voxel in the end-expiratory and other respiratory states, *m* was the total mass, and *ρ*_*end-expi*_, *ρ*_*resp*_ were the proton densities. This metric assumes conservation of mass and *S* ∝ *ρ*.

A Gaussian filter of matrix size 5x5x5 voxels was applied on the spatial domain and a 1D wrapped Gaussian filter was applied in the phase dimension for noise reduction.

### Statistical Analysis

Details of statistical analysis and image-based pulmonary function measurement are included in the supplementary material.

All scripts for this paper are available online at https://github.com/PulmonaryMRI/reproducibility.

## Results

### Representative Ventilation Maps

Figure 2 depicts the ventilation map overlayed on the registered phase-resolved lung MR images for the two registration methods and two ventilation calculation methods of one representative volunteer.

**Figure 2.**
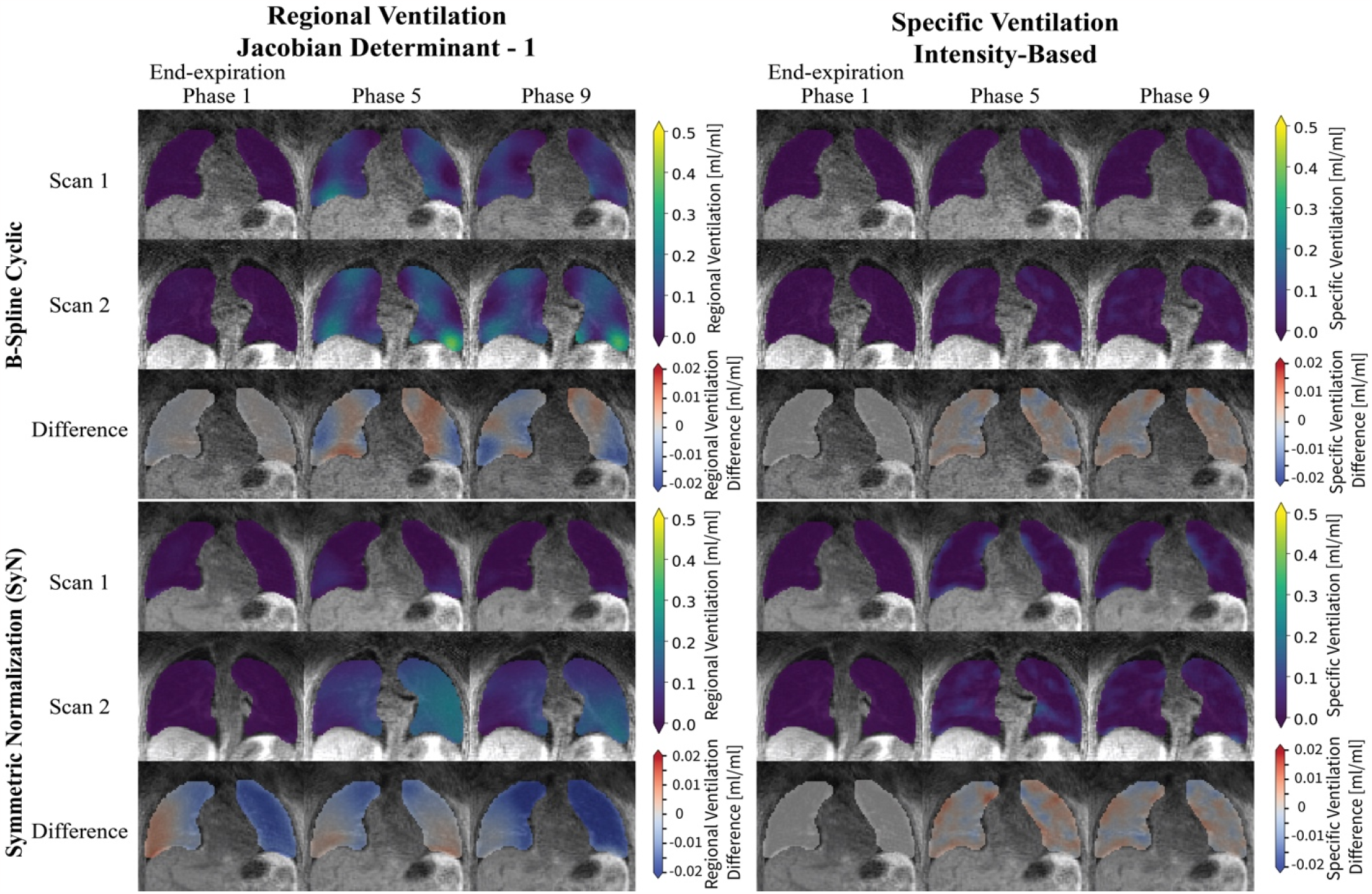
Representative Regional Ventilation Map of Two Scans and Their Difference Using Three Registration Methods. Within each method, the first row and second row are from the first and second scan respectively while the third row is the difference of ventilation maps between the two after a simple registration. A regional ventilation of 0.1 correspond to 10% of volume expansion with respect to the end-expiration state. For conciseness, three respiratory phases out of twelve are shown.

The ventilation maps for phase 1 depict the cyclic consistency of the methods. Since phase 1 is the end-expiration state and is selected to be the reference frame, the ventilation of this frame should be close or equal to 0. The intensity-based methods guaranteed it. However, for Jacobian determinant-based methods, the concatenated motion fields may not be zero if cyclic consistency is not enforced. For regional ventilation, the B-spline cyclic registration is closer to zero at phase 1, indicating that it mimics the cyclic respiratory pattern.

The regional ventilation maps show a smoother pattern compared to specific ventilation. The smoothness of regional ventilation could result from the smoothing filter applied on the motion field during registration.

### Reproducibility

#### Intra-Subject Analysis

Figure 3 shows the ventilation distributions for two representative volunteers with two registration methods and two ventilation calculation methods. This includes consistent (Volunteer 6) and inconsistent (Volunteer 5) breathing between the two scans. Results for all volunteers are shown in the supplementary Figure S1. In the split violin plots of all four combinations (columns 1-4), the median regional or specific ventilation evolves with a generally increase-decrease pattern, following the respiratory phases and matching the segmentation-based tidal volume measurements (column 5). We also computed the correlation (column 6) between lung volume measurements based on these ventilation metrics and segmented volumes across respiratory phases. For all cases, the regional ventilation approach (blue and orange) has higher Pearson correlation coefficients, r, compared to the specific ventilation approach.

**Figure 3.**
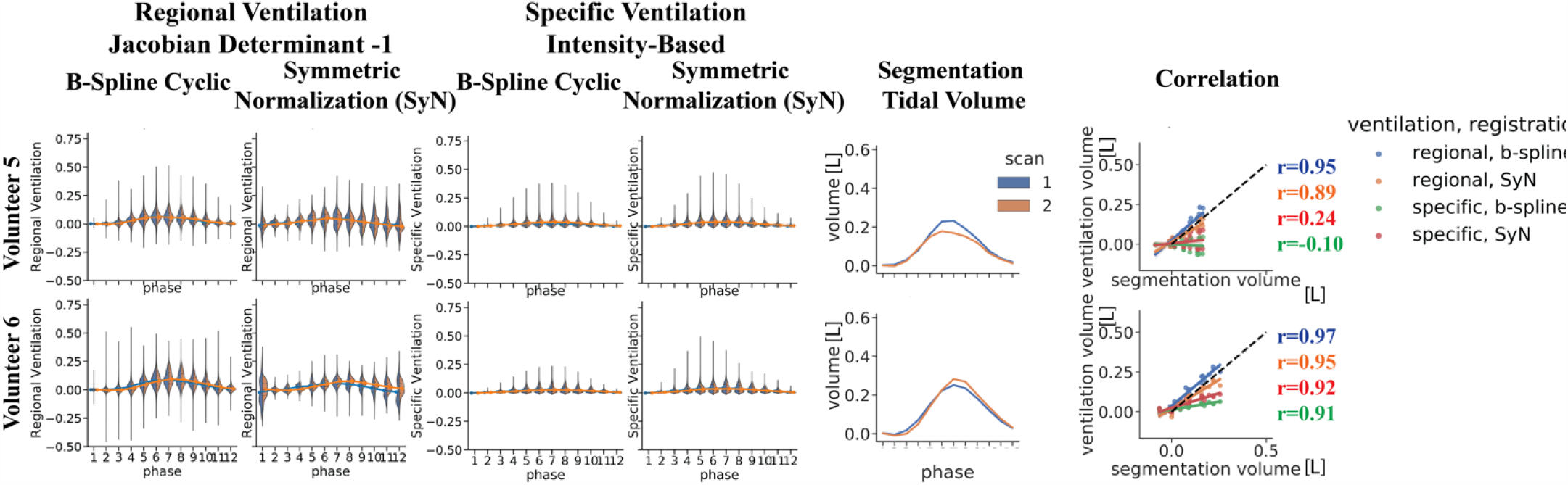
Split Violin Plot of the Ventilation Distribution, Segmentation Tidal Volume, and Their Correlation of Two Representative Subjects. The discrete horizontal axis are the respiratory phases starting from end-expiration. In the split violin plots, the two colors represent the 1^st^ and 2^nd^ scan respectively, the shaded areas depict the distribution, and the solid line across phases is the median. As for correlation, the Pearson r values are color coded for each registration and ventilation combinations. See Figure S1 for results from all volunteers.

Figure 4 compares the total ventilation as measured by each method between scans across all volunteers and respiratory phases. The Bland-Altman plots show that b-spline registration with specific ventilation has the smallest variation yet also smallest range. There are increased differences between scans as total ventilation increases, which occurs at the respiratory phases closer to end-inspiration, phases 4-8.

**Figure 4.**
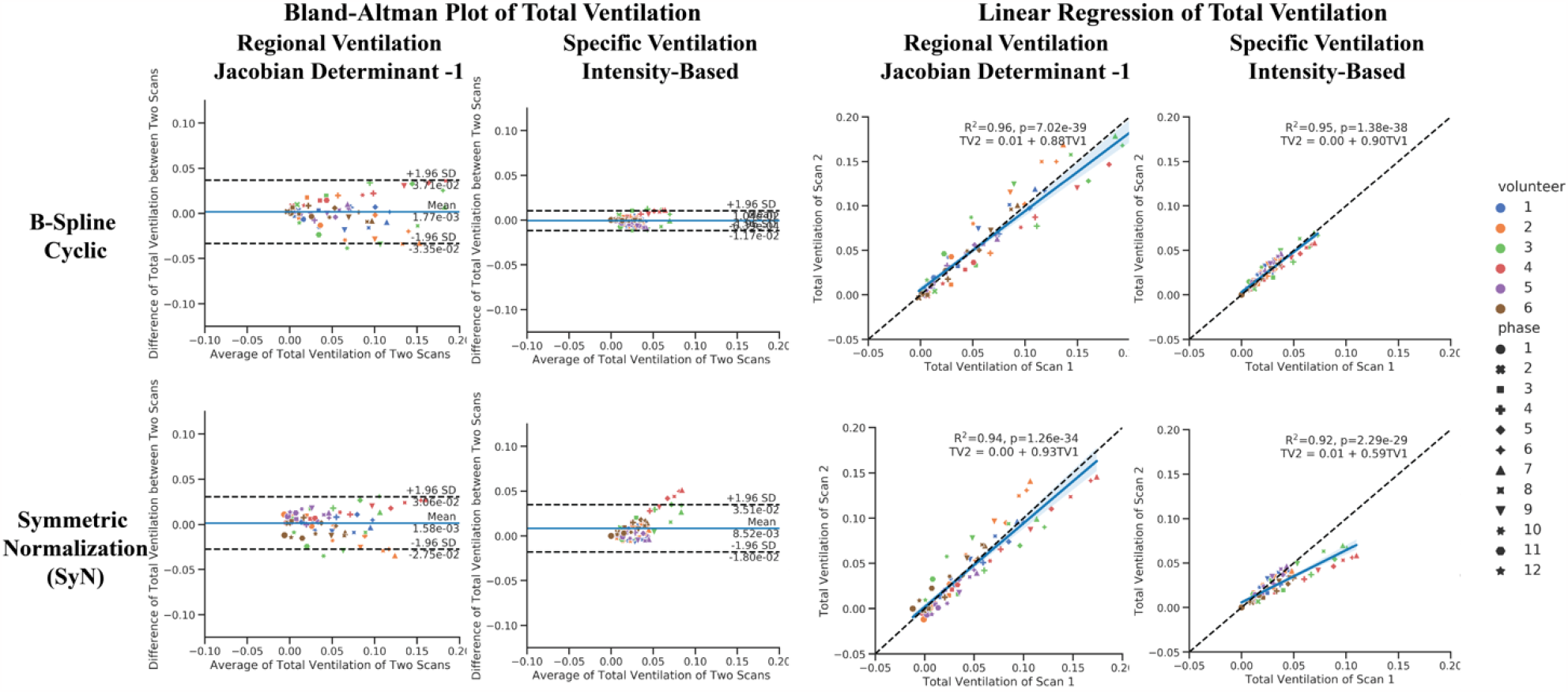
Bland-Altman Plot and Linear Regression of Total Ventilation across all subjects and respiratory phases. The total ventilation is calculated by averaging all regional ventilation within the lung volume. Each color represents a volunteer while each shape represents specific respiratory phase.

The slopes of the linear regression lines of three methods, both regional ventilation methods and b-spline registration combined with specific ventilation method, are close to 1. They also showed higher correlations, with R^2^ values 0.94-0.96. Total ventilation of Volunteers 1 and 6 align most with the dashed slope 1 line, which matches our observation from the split violin plot.

#### Sectional Ventilation

Figures 5-6 show analyses of ventilation with lungs that are separated into six different zones, lower left, lower right, middle left, middle right, upper left, and upper right sections, to learn how different sections contribute to interscan variations. Results are shown only for the cyclic b-spline registration combined with the regional ventilation approach.

**Figure 5.**
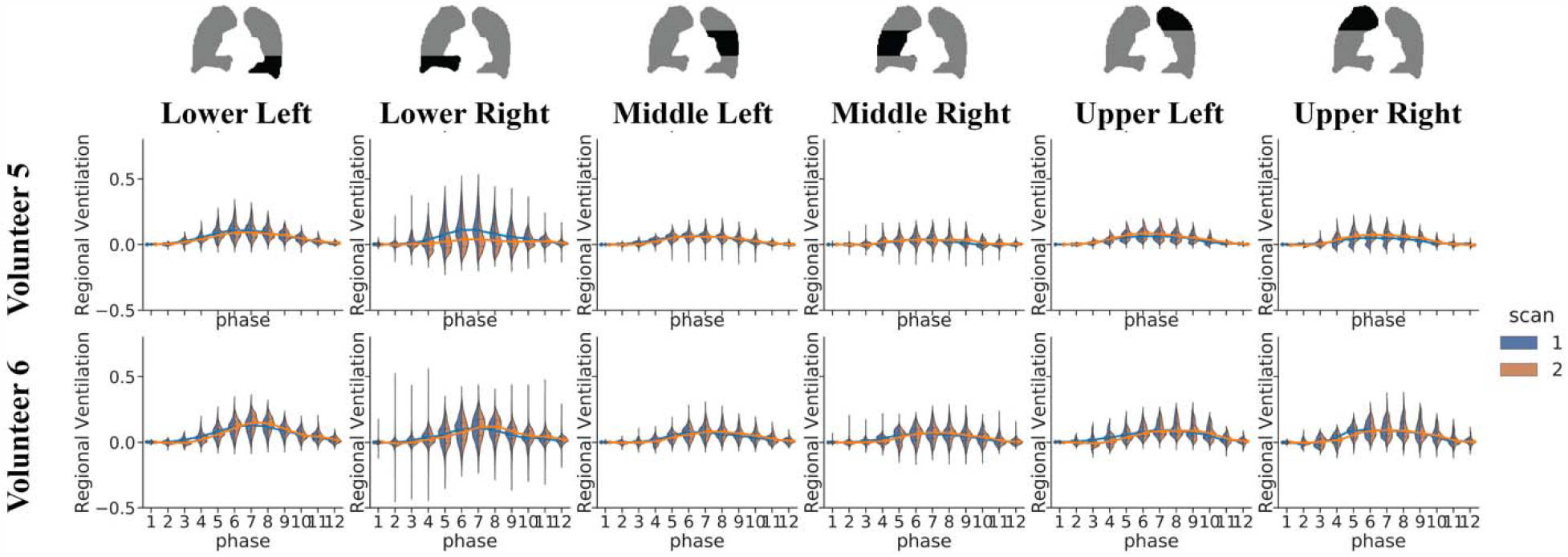
Split Violin Plots of the Ventilation Distribution for Six Lung Sections of Two Representative Subjects using the Cyclic b-Spline Registration and Regional Ventilation Method. The subjects and the violin plot characteristics are the as Figure 3. The lower left and lower right regions have the highest median ventilation among all sections. See Figure S3 for results from all volunteers.

Figure 5 and Figure S3 show that across all volunteers, there was a larger averaged ventilation value as well as a larger variance in the lower sections. This is further supported by Figure 6, where the lower left and right sections have the largest standard deviation in total ventilation difference, suggesting prominent differences and less reproducibility.

**Figure 6.**
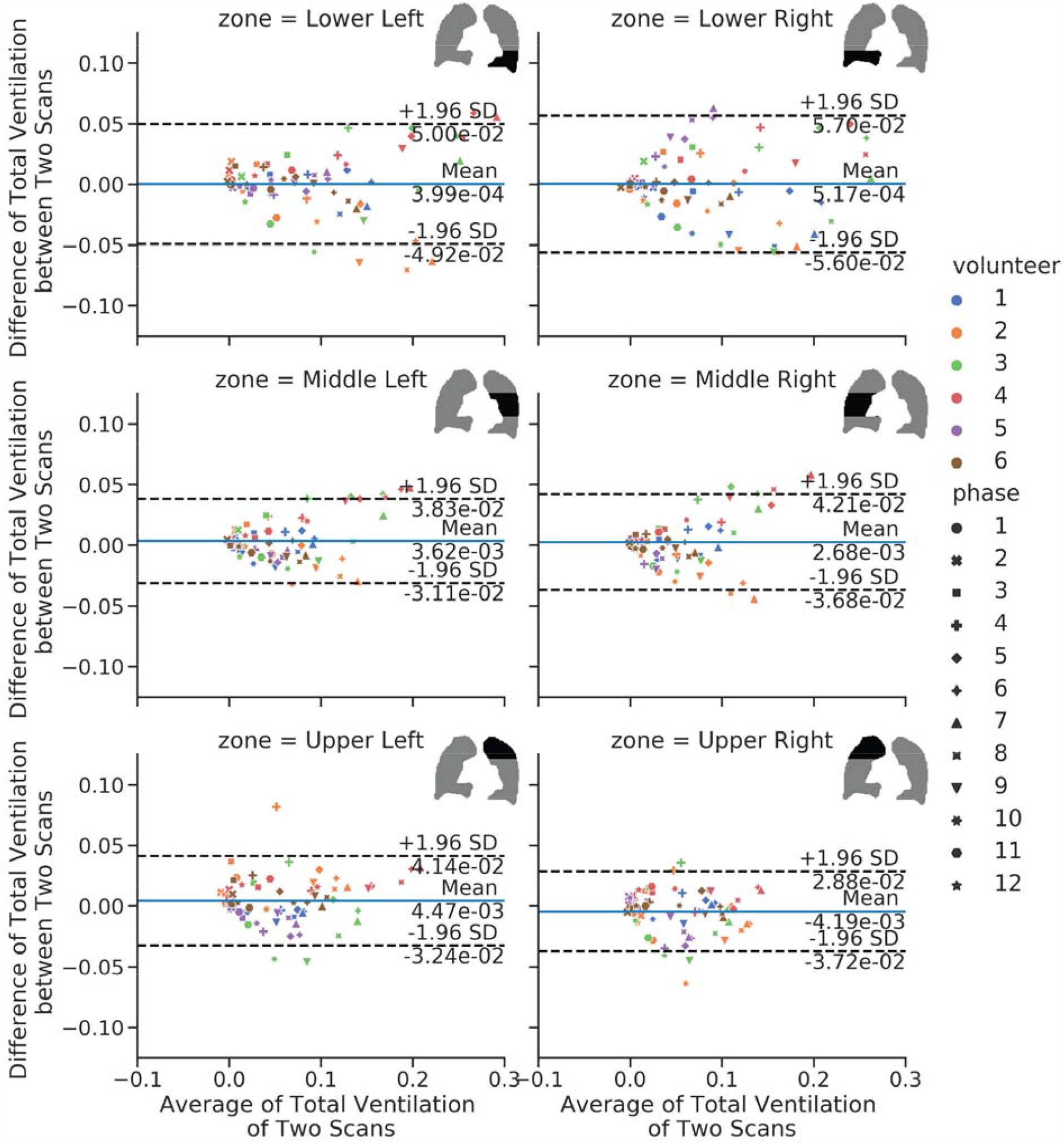
Bland-Altman Plot of Total Ventilation using the Cyclic b-Spline and Regional Ventilation Method. The lower left and lower right regions have a larger span in average total ventilation of the two scans and a larger standard deviation in the difference between the two scans.

## Discussion

In this work, we implemented a phase-resolved reconstruction for radial UTE lung MRIs and calculated the ventilation maps. We tested four approach combinations, covering two registration methods, the 3D+t b-spline cyclic registration (13) and symmetric normalization (14), and two ventilation map quantification methods, the deformation field-based regional ventilation (19,20) and the intensity difference-based specific ventilation (21,22) method. Through violin plot, coefficient of variation (Figure S2, S4), Bland-Altman plot, and linear regression, we observed different performances from these methods. The cyclic registration combined with regional ventilation has the highest correlation with segmented lung volume, while cyclic registration combined with specific ventilation was the most reproducible.

### Regional Ventilation Versus Specific Ventilation

Based on our investigation, regional ventilation correlates better with segmentation lung volume, while specific ventilation is more reproducible.

We believe several factors might contribute to this performance. Since the intensity-based specific ventilation uses the image intensity, it is more sensitive to low SNR and image registration errors particularly from partial volume effects of vessel or chest wall signals that are much higher than the lung parenchyma. Because of this, spatial and temporal averaging is required for noise reduction. This improves the reproducibility yet leads to a smaller ventilation range as shown in the split violin plots in Figure 3.

The specific ventilation calculation also relies on the assumption that the MR signal is directly proportional to proton density, but this may vary due to T2* changes (23) and the coil sensitivity map changes caused by breathing. On the contrary, the regional ventilation uses Jacobian determinant of the motion fields, which is a direct measurement of volume change ratio. This could help explain that the regional ventilation correlates better with the segmentation lung volume.

### The Effect of Registration Methods

We observed b-spline registration outperforms the SyN registration in both reproducibility and the ability to account for the cyclic nature of respiration by different registration methods.

As for cyclicity, we expect the concatenation of all motion fields from phase 1 to 12 back to phase 1 should be close to zero and thus has regional ventilation close to zero in phase 1. If we look at the ventilation colormap in the representative volunteer in

Figure 2, the regional ventilation values for phase 1 using cyclic registration are close to zero which suggests this registration is enforcing cyclic behavior. While for SyN registration, the ventilation of phase 1 has a large range, showing that cyclicity is not accounted for.

This could be explained by the algorithms of the registration methods. 3D+t b-spline registration applies b-spline grids in the wrapped temporal dimension, which assumes the first and last temporal frames are adjacent. However, in SyN registration, since the motion fields are concatenated from phase 1 to 12, slight errors in each registration step accumulate, and thus the SyN method does not account for cyclicity.

### Limitations

This study has a few limitations. First, we fixed the reconstruction to 12 respiratory phases, balancing between less streaking artifact and more respiratory phases. We speculate that with a smaller number of bins, the data to reconstruct each image will increase, leading to an increased SNR, which will potentially improve the performance of the intensity-based specific ventilation. Second, we tested one set of registration parameters on both registration methods. Second, we modified the b-spline cyclic registration parameters from (13) and the SyN registration parameters from ANTs default. The parameters were tuned by manually inspecting the quality of the registrations. However, some key parameters such as the multiscale registration levels, the grid spacing, or the smoothing factor may affect the registration performance or the smoothness of the motion field. Thus, their effects on ventilation quantification need further investigation.

## Conclusion

In this reproducibility study, we tested four combinations of registration methods and ventilation calculation methods to quantify ventilation on 3D phase-resolved ^1^H UTE MRI of healthy volunteers.

We conclude that cyclic registration is superior to SyN for ventilation purposes. The registration method significantly affects the registration-based ventilation analysis. Regional ventilation correlates better with segmentation lung volume, while specific ventilation is more reproducible.

The sectional analysis implies that the lower left and lower right sections ventilate the most during MRI scanning and account for the most between-scan ventilation differences.

## Acknowledgements

We would like to thank Dr. Kevin Johnson for generously sharing the UTE sequence, Mary Frost and Kimberly Okamoto for their assistance in the volunteer studies. We also thank the volunteers who participated in this study. This work has been supported by NIH R01HL136965.

## Tables

**Table 1.** Image-Based Pulmonary Function Measurements across all volunteers. The respiratory rate is calculated from the filtered respiratory motion curve (Figure 1B), the functional residual capacity is the segmentation volume at the end-expiration state, and the tidal volume is the segmentation volume difference of the end-inspiration and end-expiration state.

|  | Volunteer | 1 | 2 | 3 | 4 | 5 | 6 |
| --- | --- | --- | --- | --- | --- | --- | --- |
| Respiratory Rate<br>[breaths per minute] | Scan1 | 14.9±0.7 | 16.0±2.1 | 19.3±1.0 | 14.5±0.9 | 12.5±1.5 | 17.5±1.4 |
|  | Scan2 | 14.3±1.3 | 15.6±1.5 | 20.1±1.4 | 13.5±1.2 | 15.5±1.0 | 16.5±0.7 |
| Functional Residual<br>Capacity [L] | Scan1 | 3.85 | 1.59 | 1.91 | 2.14 | 3.37 | 2.77 |
|  | Scan2 | 3.60 | 1.58 | 2.00 | 2.14 | 2.75 | 2.88 |
| Tidal Volume [L] | Scan1 | 0.46 | 0.18 | 0.40 | 0.52 | 0.32 | 0.26 |
|  | Scan2 | 0.43 | 0.27 | 0.34 | 0.38 | 0.12 | 0.29 |

## Notes

This work is supported by NIH R01HL136965

### Competing Interest Statement

The authors have declared no competing interest.

