## Supplementary Material for "Pulmonary Ventilation Analysis Using ^1^H Ultra-Short Echo Time (UTE) Lung MRI: A Reproducibility Study"

**Methods**

- Statistical Analysis
  - Violin Plot

The violin plot was first introduced by Hintze and Nelson in 1998 (1). It resembles box plots but with added density of distribution to the plot. In this study, we presented the regional ventilation and specific ventilation with violin plots at all respiratory phases.

We plotted the results from two scans in different colors in a split violin plot using the seaborn toolbox (2). The quartiles are indicated by the dashed lines in the violin plot. We traced the median ventilation across respiratory phases and compared the results of the two scans with violin plots.

- Coefficient of Variation

The within-subject coefficient of variation (CV) (3) metric was used as the reproducibility measurement. Since the ventilation measurements were in percentages, the logarithm variation of CV is adopted (4). *Vent_1_* and *Vent_2_* are the ventilation measure from the 1^st^ and 2^nd^ scan, and N are the total number of voxels within the lung segmentation.

$$CV=\exp\left\{ \sqrt{\frac{1}{N}\cdot\sum_{n=1}^{N} \frac{\left[ \log\left( 1+{Vent}_{1} \right)-\log\left( 1+{Vent}_{2} \right) \right]^{2}}{2}} \right\}-1$$

- Bland-Altman

Bland-Altman plots (5,6) visualize the agreement between two repeated studies. We compared the agreement of total ventilation which is calculated by averaging the regional ventilation or specific ventilation within the lung.

We plotted the difference of total ventilation against the average total ventilation of two scans and calculated the mean and standard deviation of the differences. A mean closer to 0 and a smaller standard deviation suggests more reproducible measurements.

- Linear Regression

Linear regression fits were performed between the total ventilation calculated by average regional ventilation or specific ventilation of the first and repeated scan. R-squared, P-values, the slope and intercept were reported in the figure. The closer to a slope of 1 and an intercept of 0 suggests a more reproducible scan.

- Image-Based Pulmonary Function Measurement

Pulmonary function values of respiratory rate, vital capacity, and tidal volume were measured from the UTE images. The respiratory rate is defined as the number of breaths per minute. We used the center of the k-space self-navigator for the calculation (Figure 1b) and reported the mean and standard deviation of the number of breaths in each minute in the 8-minute tidal breathing scan.

The vital capacity and tidal volume are two parameters usually measured in the pulmonary function test. We extract them from the segmentation volume (figure 1c). The vital capacity equals the number of voxels within the lung in the end-expiration state times the voxel size. The tidal volume equals the difference in the number of voxels between end-inspiration and other respiratory states times the voxel size (2.5^3^ mm^3^).

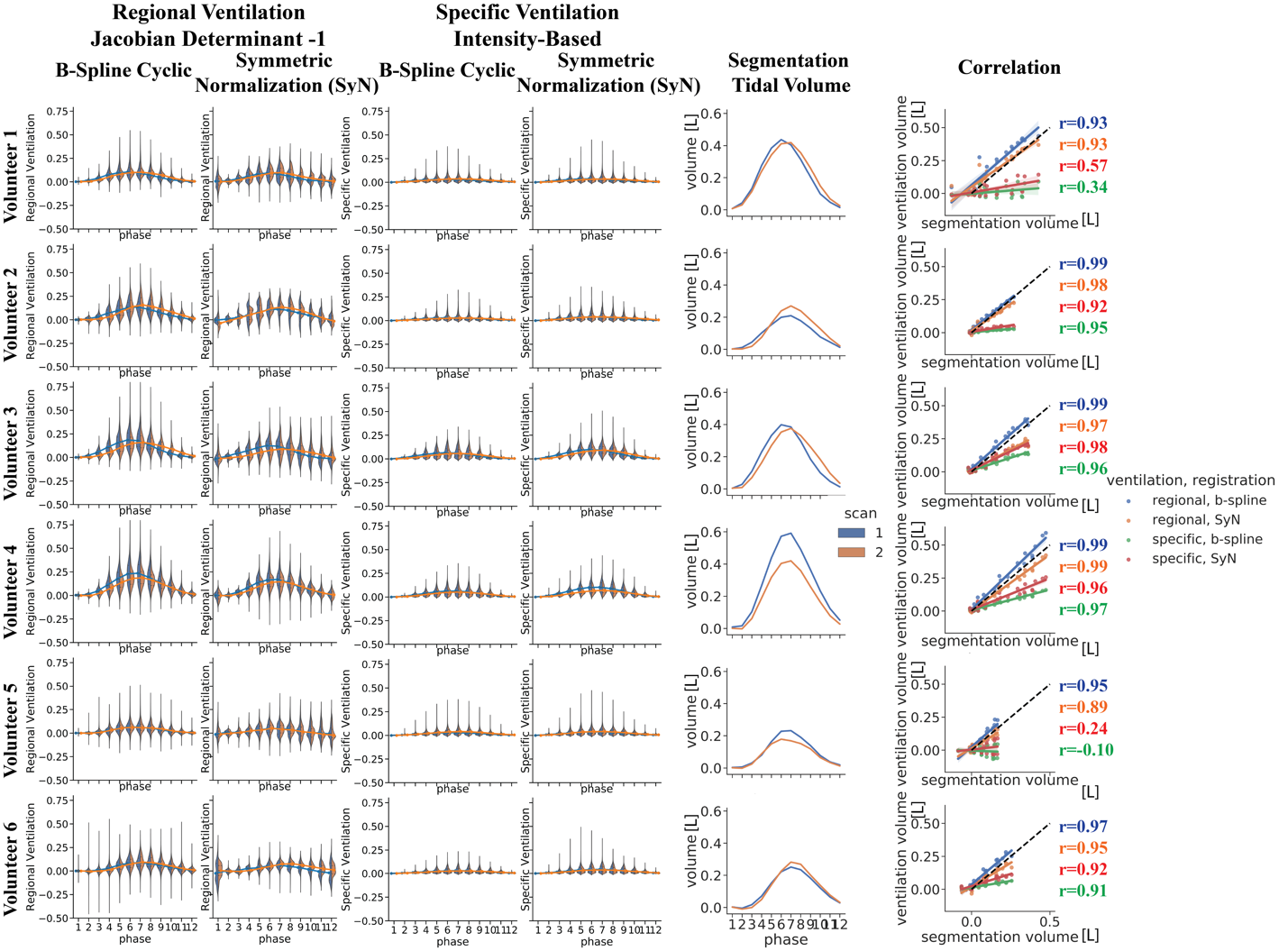

Figure S1. Split violin plots, segmentation tidal volume, and correlation between segmentation tidal volume and ventilation calculated tidal volume of all volunteers. The median regional ventilation at end-inspiration for a volunteer range from 0.15 to 0.2, corresponding to a 15%-20% volume expansion, while the median of the specific ventilation ranges from 0 to 0.1. Subjects 3 and 4 have higher ventilation in general. Subjects 1,2,3 and 6 show a high correlation between segmentation and ventilation quantification for regional ventilation methods.

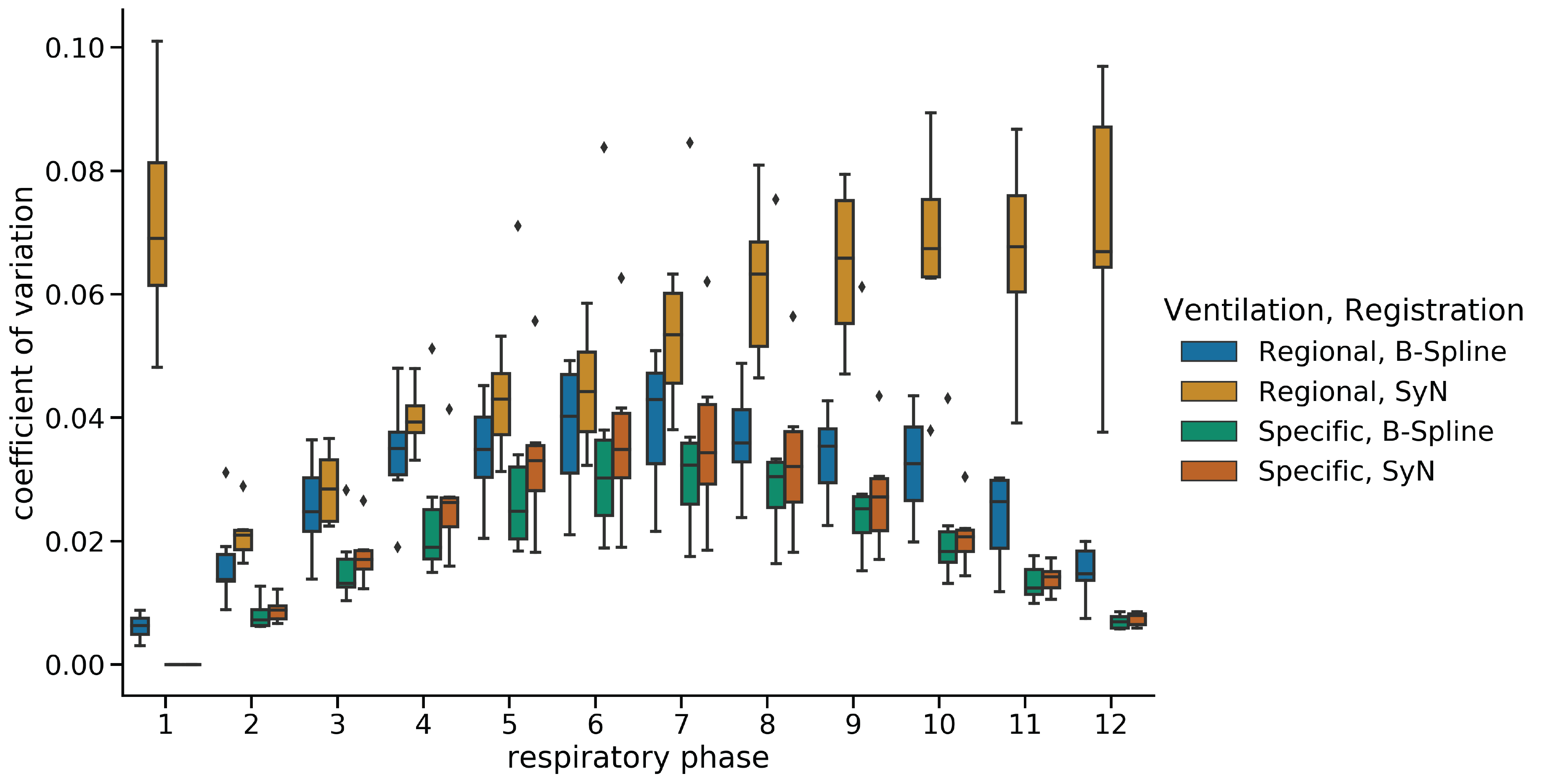

Figure S2. Within-Subject Coefficient of Variance for the ventilation methods. A value closer to zero indicates smaller differences between the two measurements, and thus more reproducible measurements. A lower coefficient of variation implies better reproducibility.

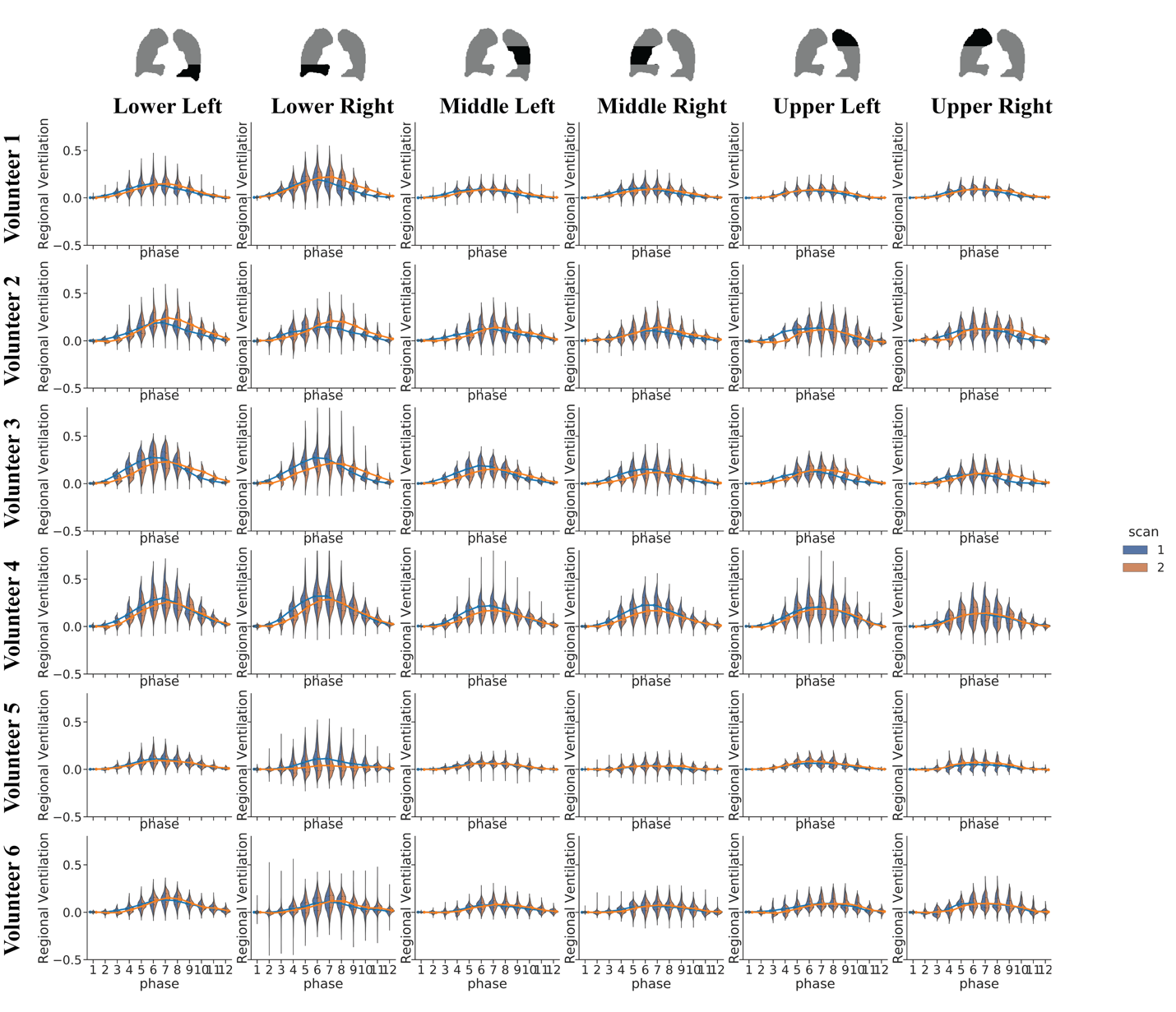

Figure S3. Split Violin Plots for Six Lung Sections for Cyclic b-Spline and Regional Ventilation of all volunteers.

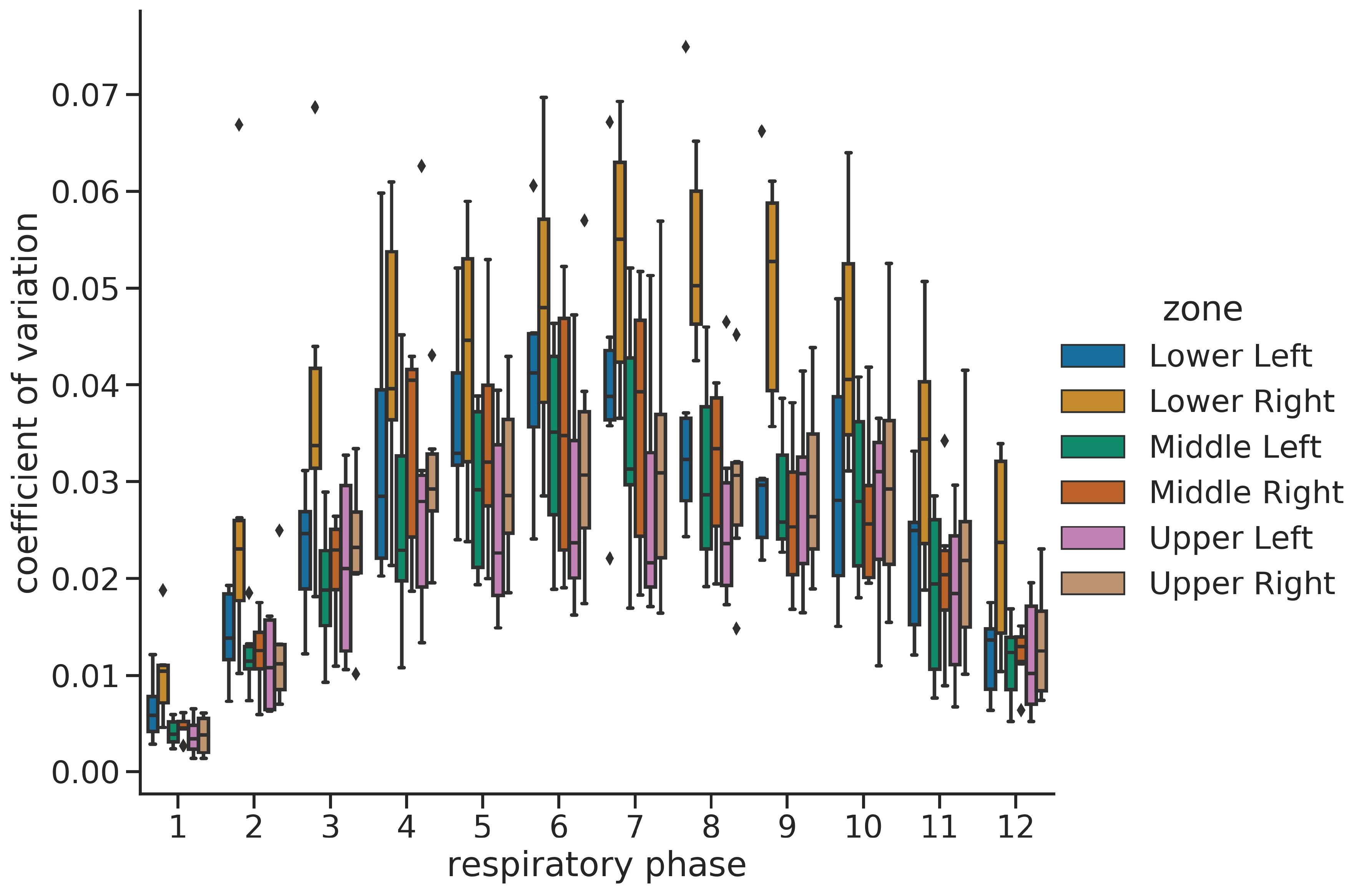

Figure S4. Coefficient of Variation (CoV) Box Plot for Six Lung Sections using the Cyclic b-Spline and Regional Ventilation Method. The lower right region has the largest CoV, suggesting it is the least reproducible, followed by the lower-left region. The other four regions have small CoVs and are similarly reproducible.
